# Chronic hM3Dq-DREADD mediated chemogenetic activation of parvalbumin-positive inhibitory interneurons in postnatal life alters anxiety and despair-like behavior in adulthood in a task and sex-dependent manner

**DOI:** 10.1101/2022.04.04.487022

**Authors:** Toshali Banerjee, Sthitapranjya Pati, Praachi Tiwari, Vidita A. Vaidya

## Abstract

Animal models of early adversity or neurodevelopmental disorders are associated with altered parvalbumin (PV)-positive inhibitory interneuron number and function, correlated with a dysregulated excitation-inhibition (E/I) balance that is implicated in the pathophysiology of neuropsychiatric disorders. We sought to address whether altering neuronal activity of PV-positive interneurons during the postnatal developmental window influences the emergence of anxio-depressive behaviors in adulthood, which are known to be perturbed in models of early adversity and neurodevelopmental disorders. We used a PV-Cre::hM3Dq-DREADD bigenic mouse line that selectively expresses the hM3Dq-DREADD receptor in PV-positive interneurons, and chemogenetically enhanced Gq signaling in PV-positive interneurons during the postnatal window via administration of the DREADD agonist, clozapine-N-oxide. Immunofluorescence studies indicated the selective expression of hM3Dq-DREADD in PV-positive interneurons in limbic circuits, and revealed a reduction in expression of the neuronal activity marker, c-Fos, in these circuits, following chemogenetic hM3Dq-DREADD mediated activation of PV-positive inhibitory interneurons. We noted no change in either growth or sensorimotor reflex milestones following chronic hM3Dq-DREADD mediated chemogenetic activation of PV-positive inhibitory interneurons in postnatal life. Adult male and female PV-Cre::hM3Dq-DREADD bigenic mice with a history of postnatal chemogenetic activation of PV-positive interneurons exhibited a reduction in anxiety and despair-like behavior in adulthood, which was noted in both a behavioral task and sex-dependent manner. These results indicate that altering neuronal activity within PV-positive interneurons during the critical postnatal developmental window can shape the emergence of anxio-depressive behaviors in adulthood, and suggest that interactions with sex as a variable play a key role in determining behavioral outcomes.

## 1 Introduction

GABAergic inhibitory pathways play a crucial role in the regulation of developmental temporal windows referred to as ‘critical periods’ (Takesian and Hensch, 2013; Reh *et al*., 2020) wherein experience exerts a profound influence on neuronal plasticity (Nabel and Morishita, 2013; Begum and Sng, 2017; Jenks *et al*., 2021), programming persistent consequences on neurocircuits and behavior (Maffei and Turrigiano, 2008; Smith-Hicks, 2013; Swanson and Maffei, 2019). Amongst the key drivers of critical period plasticity are the fast-spiking parvalbumin (PV)-positive inhibitory interneurons (Hu *et al*., 2014; Butt *et al*., 2017), that contribute to the developmental progression of the neurocircuit balance between excitation and inhibition (E/I) with a reduction in the E/I ratio associated with the closure of critical periods (Tao *et al*., 2014; Ferguson and Gao, 2018; Moore *et al*., 2018). PV-positive interneurons are present across diverse neuronal circuits, and regulate E/I balance through direct axo-somatic connections with excitatory projection neurons, driving both feed-forward and feedback inhibition. PV expression shows up within the first two postnatal weeks following birth in rodents (Seto-Ohshima *et al*., 1990; de Lecea *et al*., 1995; Fishell, 2007; Marín, 2012; Bartolini, *et al*., 2013; Wong *et al*., 2018) and matures through the early juvenile period, marked by an increase in both intensity of PV expression and the number of PV-positive interneurons, as well as the formation and strengthening of PV interneuron synapses through perineuronal net (PNN) deposition, which is associated with the eventual closure of critical windows (Favuzzi *et al*., 2017; Bucher *et al*., 2021). Indeed, it has been hypothesized that PV-positive interneuron dysregulation during these developmental windows (Marín, 2012; Bartolini *et al*., 2013; Canetta *et al*., 2016; Santiago *et al*., 2018; Murthy *et al*., 2019; Gildawie *et al*., 2020; Ohta *et al*., 2020; Gildawie *et al*., 2021; Guadagno *et al*., 2021; Klimczak *et al*., 2021; Nakamura *et al*., 2021) be one of the key pathophysiological mechanisms contributing to the neurodevelopmental origins of multiple psychiatric disorders (Ferguson and Gao, 2018; Ruden *et al*., 2021), such as major depression (Fogaça and Duman, 2019; Spijker *et al*., 2020), bipolar disorder (Konradi *et al*., 2011), autism (Contractor *et al*., 2021) and schizophrenia (Lewis *et al*., 2012; Dienel and Lewis, 2019).

PV-positive interneuron dysfunction could directly impinge on regulation of the firing of projection neurons in diverse circuits, including brain regions such as the prefrontal cortex (PFC) (Massi *et al*., 2012; Bitzenhofer *et al*., 2020) and hippocampus (Ognjanovski *et al*., 2017), that are implicated in the regulation of mood-related behavior (Ploski and Vaidya, 2021). While studies thus far have addressed the influence of modulation of PV-positive interneuron activity on anxio-depressive behaviors, most of these studies have been performed in adulthood or in the juvenile window, in brain regions such as the PFC and hippocampus (Zou *et al*., 2016; Chen *et al*., 2018; Cisneros-Franco and De Villers-Sidani, 2019; Mukherjee *et al*., 2019; Page *et al*., 2019; Nawreen *et al*., 2020; Huang *et al*., 2021; Xiao *et al*., 2021), with a paucity of information on the behavioral consequences of modulation of PV-positive interneuron activity during critical early postnatal neurodevelopmental windows (Huang *et al*., 2021; Medendorp *et al*., 2021). While animal models of early adversity and genetic models of schizophrenia and autism spectrum disorder exhibit in common a disruption of mood-related behavior, PV interneuron number and function (Ruden *et al*., 2021), it remains unclear whether directly modulating PV-positive interneuron activity during critical developmental windows can influence the emergence of anxio-depressive behaviors in adulthood.

Here, we have addressed the consequences of a chronic postnatal activation of PV-positive inhibitory interneurons on the emergence of adult anxio-depressive behaviors. We used a strategy of chemogenetically activating PV-positive interneurons in bigenic mouse models (PV-Cre::hM3Dq-DREADD) (Panthi and Leitch, 2021) expressing the hM3Dq-DREADD under the control of the PV-promoter, and stimulated the hM3Dq-DREADD with the exogenous ligand clozapine-N-oxide (CNO) in bigenic mouse pups during the postnatal window (postnatal day: P2 -P14). Our results indicate that chemogenetic PV-positive interneuron activation during this postnatal window results in a significant decline in the emergence of adult anxiety and despair-like behavior, in a task-specific and sex-dependent manner.

## 2 Materials and Methods

### 2.1 Animals

Bigenic PV-Cre::hM3Dq-DREADD mice were generated by crossing females from the PV-Cre mouse line (B6.129P2-Pvalbtm1(cre)Arbr/J; Jax stock 008069; Jackson Laboratories, USA) (Hippenmeyer *et al*., 2005) with male floxed hM3Dq-DREADD mice (B6N;129-Tg(CAG-CHRM3*-,mCitrine)1Ute/J); Jax stock 026220; Jackson Laboratories) (Zhu *et al*., 2016), and were maintained in the Tata Institute of Fundamental Research (TIFR) animal facility. The PV-Cre mouse line was maintained on a C57BL/6J background, while the floxed hM3Dq-DREADD mouse line was maintained on a mixed background of 129×1/SvJ and 129S1/Sv. All animals were kept on a twelve hour light-dark cycle (7:00 AM to 7:00PM) with *ad libitum* access to food and water. Animal genotypes were determined using PCR-based analysis with published primers available on the Jackson Laboratories website. All experimental procedures were carried out in accordance with the guidelines of the Committee for the Purpose of Control and Supervision of Experiments on Animals (CPCSEA), Government of India and were approved by the Institutional Animal Ethics Committee at TIFR.

### 2.2 Drug treatment paradigms

Bigenic PV-Cre::hM3Dq-DREADD mice pregnant dams were separated and housed individually 2-3 days before parturition. Ample nesting material in the form of paper shavings were provided across all cages. Pups born on the same day were mixed across litters and distributed across each dam to reduce litter-based variations and to keep litter size constant (8 pups per litter). The DREADD agonist, clozapine-N-oxide (CNO; Cat no. 4936; Tocris, UK) was used to selectively stimulate the excitatory DREADD, hM3Dq. Litters were divided at random into vehicle and postnatal CNO (PNCNO) treatment groups (8 litters per group). The litters were weighed every day from postnatal day 2 to 14, and the average weight of each pup in the litter was calculated to estimate the amount to be administered. Pups were orally administered CNO (1 mg/kg) or vehicle (5% sucrose in water; 1 mg/ml (w/v)) once daily from P2 - P14 (Pati *et al*., 2020). During the postnatal treatment, the pups across representative litters were assayed for the development of reflex behaviors. Between postnatal days 21-25, the pups were weaned from their dams and group-housed in same sex cohorts. The animals were left undisturbed till 2-3 months of age, following which both male and female vehicle and PNCNO-treated bigenic PV-Cre::hM3Dq-DREADD mice were assessed on behavioral tasks for anxiety and despair-like behaviors.

### 2.3 Western blotting

Samples for PNCNO experiments were generated by feeding vehicle or CNO (1 mg/kg) to PV-Cre::hM3Dq-DREADD animals from P2 - P7. Thirty minutes post vehicle or CNO treatment on P7, pups were anaesthetized on ice and sacrificed. Cortices and hippocampi were dissected and snap frozen in liquid nitrogen. The samples were subsequently stored at -80°C till further processing. The dissected tissue was homogenized using plastic dowels in ice-cold Radioimmunoprecipitation Assay (RIPA) Buffer (10mM Tris-Cl (pH 8.0), 1mM EDTA, 1% Triton X-100, 0.1% Sodium deoxycholate, 0.1% SDS, 140mM NaCl), supplemented with protease and phosphatase inhibitors (Sigma-Aldrich, USA). Protein concentrations were estimated using a Quantipro BCA assay kit (Sigma-Aldrich, USA). Protein lysates were prepared in a mixture of RIPA and Lamelli Buffer to produce a final protein concentration of 5 ug/ul. To detect the expression of the HA-tagged hM3Dq-DREADD and c-Fos, 50 ug of protein lysate was loaded onto 10% sodium dodecyl sulphate polyacrylamide electrophoresis (SDS-PAGE) gels. Protein lysates run on SDS-PAGE gels were transferred onto polyvinylidene fluoride (PVDF) membranes, and blots were incubated in 5% BSA (Sigma-Aldrich, USA) made in 1X TBST for two hours at room temperature, followed by an overnight incubation at 4°C with rabbit anti-HA antibody (1:1000, Cat no. H6908, Sigma-Aldrich, USA). For c-Fos, blots were incubated for two hours in 5% milk in 1X TBST for blocking followed by an overnight incubation with rabbit anti-c-Fos antibody (1:1000, Cat. No. 2250, Cell Signaling Technology, USA) at 4°C. Blots were also incubated with rabbit anti-actin antibody (1:1000, Cat. No. AC026, Abclonal Technology, USA) as a loading control for normalization. Blots were then subjected to serial washes followed by incubation with the secondary antibody, HRP-conjugated goat anti-rabbit (1:6000; Cat. No. AS014, Abclonal Technology) for one and a half hours at room temperature. A western blotting detection kit (WesternBright ECL, Advansta, United States) was used to detect chemiluminescent signal which was visualized using the GE Amersham Imager 600 (GE life sciences, United States). Densitometric analysis was carried out using Fiji software.

### 2.4 Immunofluorescence analysis

Coronal brain sections (P14: 60 µm) were generated on a vibratome (Leica, Germany) from postnatal (P14) PV-Cre::hM3Dq-DREADD bigenic mice. Sections were subjected to blocking in 0.1M phosphate buffer with 0.3% Triton X-100 (PBTx) containing 10% horse serum (Thermo Fisher Scientific, Cat. No. 26-050-088) and 0.5% bovine serum albumin (Sigma-Aldrich) for two hours at room temperature. Double-label immunofluorescence experiments were conducted to assess (1) the colocalization of the HA-tagged hM3Dq-DREADD or (2) the neuronal activity market c-Fos with PV-positive inhibitory interneurons. Coronal sections containing the medial prefrontal cortex (mPFC) or hippocampus were incubated with (1) an antibody cocktail of rat anti-HA (1:200, Cat. No. 10145700, Roche Diagnostics, USA) with rabbit anti-PV (1:500, Abcam, Cat. No. ab11427, USA) for three days at room temperature; or (2) with a cocktail of rabbit anti c-Fos (1:500, Cell Signaling Technology, Cat. No. 2250) and mouse anti PV (1:500, Sigma, Cat. No. SAB4200545) for one day at room temperature. Sections were then subjected to serial washes in 0.1M phosphate buffer for fifteen minutes each prior to incubation with secondary antibodies, for two hours at room temperature: (1) goat anti-rabbit IgG conjugated to Alexa Fluor 568 (1:500; Cat. No. A-11011, Invitrogen, USA) and goat anti-rat IgG conjugated to Alexa Fluor 488 (1:500; Cat. No. A-21212, Invitrogen) to visualize PV - HA double immunofluorescence and (2) anti-rabbit IgG conjugated to Alexa Fluor 555 (1:500, Cat. No. A-31572) and anti-mouse IgG conjugated to Alexa Fluor 488 (1:500, Cat. No. A-21202) to visualize PV - c-Fos double immunofluorescence. Sections were washed in 0.1M phosphate buffer and mounted on slides using Vectashield Antifade Mounting Medium with DAPI (H-1200, Vector Laboratories, USA) and images were visualized on a FV1200 confocal microscope (Olympus, Japan).

### 2.5 Behavioral Assays

#### 2.5.1 Postnatal Reflex Behaviors

During the PNCNO treatment, pups were tested on P7, P10 and P13 to assess the ontogenic development of reflex behaviors, to determine whether CNO administration during this developmental window influences reflex behaviors. The development of the following reflexes were assessed in both vehicle and PNCNO-treated PV-Cre::hM3Dq-DREADD bigenic mouse pups, namely: a) surface rightings, which assessed the time taken in seconds for a pup that was placed inverted to realign itself onto its paws, b) negative geotaxis, wherein pups were placed at a 30 degree angle with the head pointing downwards, and the time taken in seconds for the pup to reorient itself with the head pointed upwards was determined, c) air righting, where pups were dropped into their home cage from a height of 25 cm while upside-down and the ratio of the successful attempts to land on its paws in ten such events was calculated. Reflex behaviors were assessed across pups (n = 24) derived from 3-4 litters/treatment group.

#### 2.5.2 Adult Behaviors

PNCNO-treated PV-Cre::hM3Dq-DREADD mice and their respective vehicle-treated cohorts were subjected to behavioral tests to assess anxiety and despair-like behaviors in adulthood. Given the large cohorts to be processed for behavioral analysis, we segregated the adult vehicle and PNCNO-treated PV-Cre::hM3Dq-DREADD male and female cohorts, and ran both sexes through behavioral analyses separately. For anxiety-like behaviors, animals were assessed on the open field test (OFT), elevated plus maze (EPM) test and light-dark avoidance test (LD Box) and for despair-like behaviors, animals were subjected to the tail suspension test (TST) and forced swim test (FST). All tests were recorded using Ethovision X11(Noldus, Netherlands) and analyzed using its automated tracking software or manually (for TST and FST). The OFT, EPM and LD Box arenas were cleaned with 80% ethanol in between trials. For the FST, the tank water was changed every four trials. A wash-out of 7-10 days was allowed between each behavioral test during which the animals were left undisturbed in their home cage except for standard animal facility handling.

#### Open Field test (OFT)

The OFT arena was a 40 cm x 40 cm x 40 cm wooden box, with a center area of 20 cm x 20 cm. Animals were introduced to the arena by being placed at any of the four corners of the box, facing towards the center. The behavior was recorded for 10 minutes after placing the animal in the arena, using an overhead analog camera (Harvard Apparatus, United States). Ethovision X11 was used to track the behavior. Parameters such as total distance travelled in arena, percent distance travelled in the center of the arena and percent time spent in the center of the arena were determined.

#### Elevated Plus Maze test (EPM)

The EPM arena was an elevated 4-armed cross maze with alternating open and closed arms at a height of 50 cm above the ground, with each arm having an area of 30 cm x 5 cm. The walls of the closed arms were 15 cm in height. Animals were introduced to the 5 cm x 5 cm center, facing the intersection between a closed arm and open arm. The behavior was recorded for 10 mins after introducing the animal to the arena, using the overhead camera and analyzed using Ethovision X11. Parameters such as total distance travelled, distance travelled in open and closed arms, percent distance travelled in open and closed arms, percent time spent in open arms and number of entries into open arms were analyzed.

#### Light-Dark Avoidance Test

The Light-Dark Box arena consisted of a 25 × 25 cm light and a 15 × 25 cm dark chamber, connected by a 10 × 10 cm opening. Animals were introduced at the light-dark interface, facing the light chamber. Behavior was recorded for ten minutes using an overhead camera and the videos were manually analyzed by an experimenter blinded to the treatment groups. Parameters such as number of entries in the light chamber and percent time spent in the light chamber were determined.

#### Tail Suspension test (TST)

Animals were suspended by their tail at a height of 50 cm above the ground, and behavior was recorded for six minutes using a Logitech C250 webcam aligned parallel to the behavioral set-up. Behavior was analyzed for the last five minutes by an experimenter blinded to the treatment conditions. Immobility was determined as an event when the animal lies still, without any active struggle. Number of immobility events and the percent time spent immobile was assessed.

#### Forced Swim test (FST)

The FST tank was a plexiglass cylinder, 50 cm tall with an inner diameter of 14 cm. Water at room temperature (25-29 □) was added to each cylinder to a height of 30 cm. Animals were placed in the tank and behavior was recorded for six minutes using a Logitech C250 webcam aligned parallel to the tanks. Immobility was determined as an event characterized by the animals floating passively at the surface of the water, without any active struggle. Number of immobility events and percent time spent immobile was quantified by an experimenter blinded to treatment conditions, for the last five minutes of the recording.

### 2.6 Statistical analysis

Changes in weight profile and reflex behaviors upon PNCNO and vehicle administration to bigenic PV-Cre::hM3Dq-DREADD animals were subjected to a two-way ANOVA with repeated measures. Experimental results from western blot analysis and all behavioral measures were subjected to a two-tailed, unpaired Student’s *t*-test with Welch’s correction applied in cases of unequal variance across control and treatment groups. The Kolmogorov-Smirnov test was used to determine normality. All statistical analysis and graphing was done using GraphPad Prism (GraphPad Software Inc, USA). Data are expressed as mean ± standard error of mean (S.E.M) with statistical significance set at *p* < 0.05.

## 3 Results

### 3.1 Expression and activation of hM3Dq-DREADD in parvalbumin-positive inhibitory neurons in PV-Cre::hM3Dq-DREADD bigenic mice in the postnatal window

We generated PV-Cre::hM3Dq-DREADD bigenic mice (Figure 1A) to examine the influence of hM3Dq-DREADD mediated activation of PV-positive inhibitory interneurons during the critical postnatal temporal window (P2 - P14) on the emergence of anxio-depressive behaviors in adulthood. We first examined the expression of the HA-tagged hM3Dq-DREADD in postnatal pups at P14, and noted robust HA expression in both the medial prefrontal cortex (mPFC, Figure 1B-E) and hippocampus (dorsal CA1 and dorsal DG, Figure 1F-M) using double immunofluorescence analysis. We further ascertained the expression of the HA-tagged hM3Dq-DREADD receptor at P7 (Figure 1N) using western blotting analysis, which revealed robust HA expression in the cortex and hippocampus (Figure 1O). We sought to examine whether chronic CNO-mediated hM3Dq-DREADD activation of PV-positive interneurons altered neuronal activity in the cortex and hippocampus in postnatal PV-Cre::hM3Dq-DREADD pups that received daily administration of CNO (1 mg/kg) or vehicle treatment from P2-P7 via assessing expression of the neuronal activity marker, c-Fos (Figure 1P). Quantitative densitometric analysis of c-Fos protein levels indicated a significant decline in c-Fos protein expression in the hippocampus (*p* = 0.03), and a trend towards a significance (*p* = 0.1) for the decline in c-Fos protein levels in the cortex (Figure 1Q) of CNO administered PV-Cre::hM3Dq-DREADD pups at P7, as compared to their age-matched vehicle-treated controls. We also administered an acute treatment of CNO or vehicle to bigenic PV-Cre::hM3Dq-DREADD pups at P14 to assess the expression of the neuronal activity marker, c-Fos in PV-positive interneurons using double immunofluorescence. We noted an induction of c-Fos in PV-positive interneurons, as revealed in representative images from the cortex of acute CNO and vehicle-treated pups (Figure 1R). Taken together, these results indicate that the HA-tagged hM3Dq-DREADD is expressed in PV-positive interneurons during the postnatal window based on both immunofluorescence and western blotting analysis, and that chronic CNO treatment (P2 - P7) to PV-Cre::hM3Dq-DREADD bigenic mice evokes a overall reduction in c-Fos protein levels in the hippocampus as revealed by densitometric analysis.

**Figure 1.**
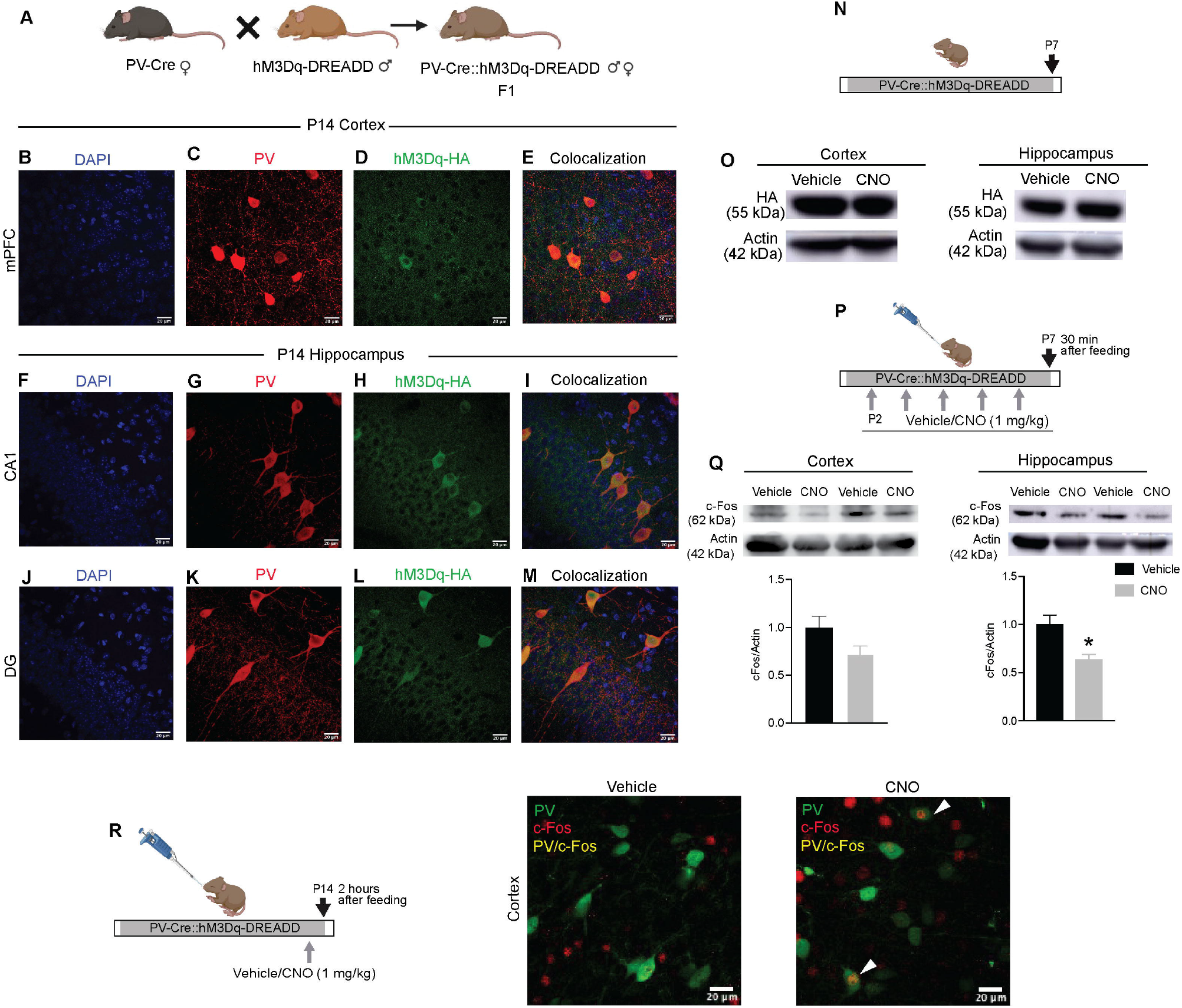
Expression and activation of hM3Dq-DREADD in parvalbumin-positive inhibitory interneurons in PV-Cre::hM3Dq-DREADD bigenic mice in the postnatal window (A) Shown is a schematic representation of female parvalbumin (PV-Cre) mice crossed with male hM3Dq-DREADD mice to create F1 progeny of bigenic PV-Cre::hM3Dq-DREADD mice that selectively express the hM3Dq-DREADD in PV-positive inhibitory interneurons. Shown are representative immunofluorescence images taken for postnatal day 14 pups (P14, n = 3 - 4) indicating the colocalization of PV-positive neurons (red) with the HA-tagged hM3Dq-DREADD (green) in the mPFC (B-E), CA1 (F-I) and dentate gyrus (DG) (J-M). Shown is a schematic of the P7 bigenic PV-Cre::hM3Dq-DREADD mouse pups sacrificed for western blotting analysis in the cortex and hippocampus to detect the HA-tagged hM3Dq-DREADD (N). Shown are representative immunoblots of HA, and the associated actin loading controls, that indicate the expression of HA-tagged hM3Dq DREADD receptors in the cortex and hippocampus of P7 bigenic PV Cre::hM3Dq-DREADD mouse pups (O). Shown is the schematic representation of the treatment paradigm of bigenic PV Cre::hM3Dq-DREADD mouse pups with vehicle or CNO (1 mg/kg) from P2 - P7 to evoke hM3Dq-DREADD-mediated activation of PV-positive neurons, sacrificed thirty minutes after the last vehicle/CNO treatment (P). (Q) Shown are representative immunoblots and densitometric quantitation for c-Fos expression, normalized to the loading controls actin, reveal a trend decline in c-Fos protein levels in the cortex, and a significant decline in c-Fos protein expression in the hippocampus in the CNO-treated group. (R) Shown is a schematic treatment paradigm wherein bigenic PV Cre::hM3Dq-DREADD mouse pups were treated with vehicle or CNO (1 mg/kg) at P14 to activate the hM3Dq-DREADD in PV-positive interneurons, and sacrificed two hours after the treatment. Representative immunofluorescence images reveal an induction of c-Fos (red) in PV-positive interneurons (green) in the cortex of CNO treated postnatal day 14 pups (P14, n = 3 per group) in comparison to their vehicle treated controls, indicating an hM3Dq-DREADD mediated activation of PV-positive inhibitory interneurons. Results are expressed as the fold change of vehicle and are the mean ± S.E.M. (\**p* < 0.05 as compared to their vehicle treated controls, n = 3 - 4 per group, Student’s *t*-test).

### 3.2 Chronic chemogenetic activation of PV-positive inhibitory neurons during the early postnatal window does not alter body weight or the normal ontogeny of reflex behaviors

We examined the consequences of hM3Dq DREADD mediated activation of PV-positive inhibitory neurons during the early postnatal window (P2 - P14) on body weight and on the emergence of sensorimotor developmental milestones. Bigenic PV-Cre::hM3Dq-DREADD mouse pups received either the DREADD agonist, CNO (1 mg/kg) or vehicle once daily from P2 to P14 (Figure 2A). Analysis of the average weights of pups across eight litters each from the vehicle and postnatal CNO (PNCNO) treated PV-Cre::hM3Dq-DREADD bigenic mouse pups were recorded daily from P2 - P14. Repeated measures ANOVA analysis revealed that while there was a significant weight gain (F_(12, 168)_ = 129.4; *p* < 0.0001) across all pups as expected across this time window (P2 - P14), this did not differ across the vehicle and PNCNO-treated mouse pups, indicating that PNCNO administration does not alter normal growth milestones. We next examined the emergence of sensorimotor reflex behaviors (Figure 2C-E) which were assessed on P7, P10 and P13, to examine whether CNO administration impacts the ontogeny of reflex behaviors. Two-way ANOVA analysis for surface rightings (Figure 2C) showed no significant difference in time taken for realignment between treatment groups, with both groups showing a significant decline in time taken for successful righting across the developmental time window (F _(2, 134)_ = 25.50; *p* < 0.0001). Two-way ANOVA analysis for negative geotaxis (Figure 2D) revealed no significant difference in time taken for reorientation, across treatment groups and across the developmental time window. Two-way ANOVA analysis for air rightings (Figure 2E) indicated no significant difference in ratio of successful realignments, between treatment groups, with both groups showing a significant improvement in successful realignments across the developmental time window examined (P7-13) (F _(2, 134)_ = 143.1; *p* < 0.0001). Collectively, these results indicate that postnatal hM3Dq-DREADD mediated chemogenetic activation of PV-positive interneurons does not alter normal weight developmental milestones during postnatal life, and does not influence the ontogeny of sensorimotor reflexes.

**Figure 2.**
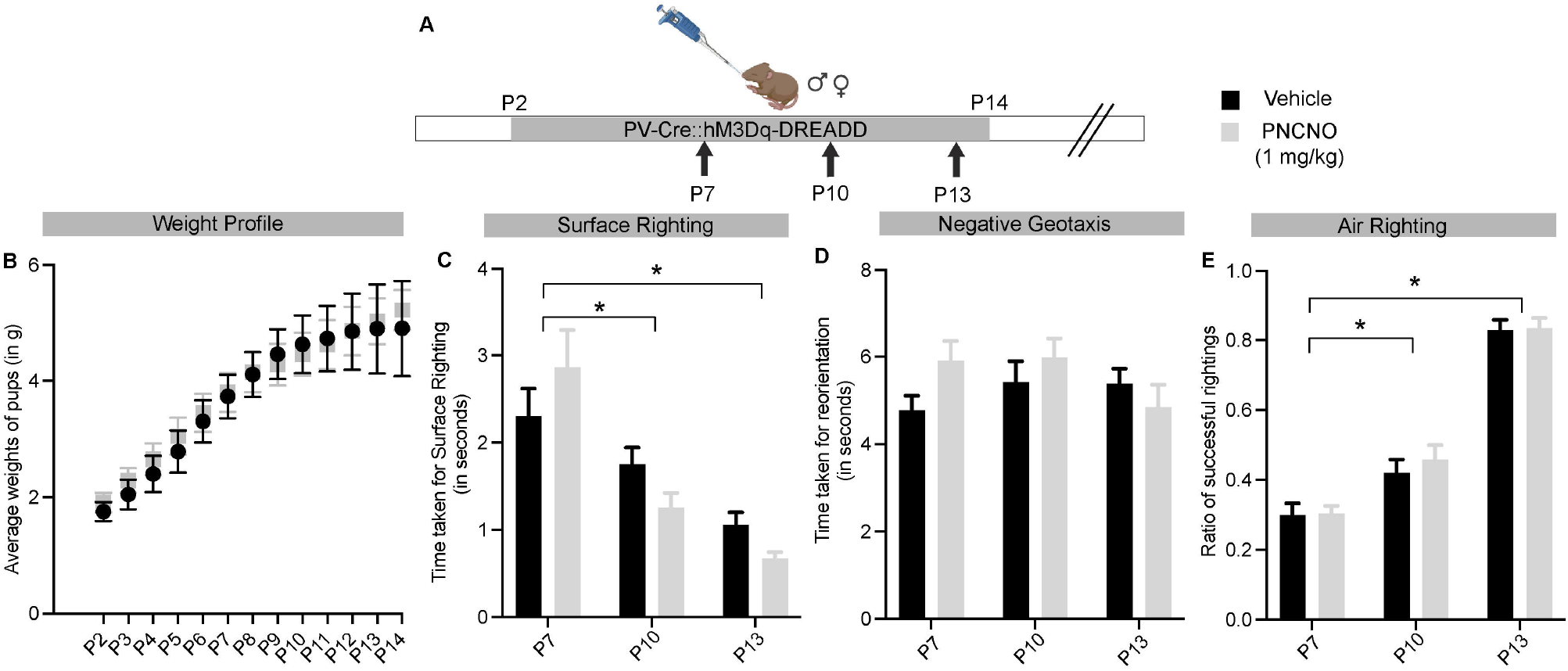
Chronic chemogenetic activation of PV-positive inhibitory interneurons during the early postnatal window does not alter body weight or the normal ontogeny of reflex behaviors (A) Shown is a schematic representation of the treatment paradigm of bigenic PV Cre::hM3Dq-DREADD mouse pups, male and female, with vehicle or CNO (1 mg/kg) from P2 - P14 to evoke hM3Dq-DREADD-mediated activation of PV-positive neurons, with weight profiling carried out daily and surface righting, negative geotaxis and air-righting reflexes profiled at P7, P10 and P13. (B) Shown is the weight profile of vehicle and PNCNO-treated bigenic PV Cre::hM3Dq-DREADD mouse pups measured daily from P2-14 indicating no change in gross weight during the treatment, with both groups demonstrating a similar age-dependent increase in weight. Profiling of the ontogeny of reflex development indicated no change between the vehicle and PNCNO-treated bigenic PV-Cre::hM3Dq-DREADD mouse pups in time taken for surface righting (C), time taken for reorientation when profiling negative geotaxis (D) and ratio of successful rightings when profiling the air-righting reflexes (E) at P7, P10 and P13. The normal progression of reflex development indicated a significant reduction in time taken for surface rightings on P10 and P13 as compared to P7 in both treatment groups (C). The ratio of successful rightings also significantly increased with developmental age in P10 and P13 time-points as compared to P7, in both treatment groups (E). Results are expressed as the mean + S.E.M. (\**p* < 0.05 as compared to their vehicle treated controls, Main effect of age, Two-way ANOVA analysis; n = 24 per group).

### 3.3 Chronic chemogenetic activation of PV-positive inhibitory interneurons during postnatal life evokes persistent changes in anxiety-like behavior in adult male and female mice in a task-specific manner

Adult male and female PV-Cre::hM3Dq-DREADD mice that received vehicle or PNCNO treatment from postnatal day P2 - P14 were subjected to behavioral analysis for anxiety-like behavior on the OFT, EPM and LD box in adulthood (Figure 3A). In the OFT, PNCNO-treated bigenic PV-Cre::hM3Dq-DREADD males, as compared to their vehicle-treated controls, showed a significant increase in the total distance travelled in the arena (Figure 3C, *p* = 0.044) and the number of entries to the center of the arena (Figure 3F, *p* = 0.0485). We noted no significant difference between the treatment groups with regards to measures such as percent distance travelled in the center of the OFT arena (Figure 3D) and percent time spent in the center of the arena (Figure 3E). In the EPM test, PNCNO-treated bigenic PV-Cre::hM3Dq-DREADD males exhibited no significant differences in the total distance travelled in the maze (Figure 3H). We observed a significant increase in the percent distance traversed in the open arms of the EPM (Figure 3I, *p* = 0.011), accompanied by a significant increase in the number of entries into the open arms (Figure 3K, *p* = 0.0002) and a trend towards significance for the percent time spent in the open arms (Figure 3J, *p* = 0.09). We also noted a significant decline in both the percent distance travelled in the closed arms (Figure 3L, *p* = 0.002) and the percent time spent in the closed arms of the EPM (Figure 3M, *p* = 0.017). We did not observe any difference between the treatment groups for the number of entries to the closed arms (Figure 3N). PNCNO-treated bigenic PV-Cre::hM3Dq-DREADD males did not differ from the vehicle-treated control group in their behavior on the LD box in adulthood, with no differences noted across the treatment groups on measures such as the number of entries into the light chamber (Figure 3P) and the percent time spent in the light chamber (Figure 3Q).

**Figure 3.**
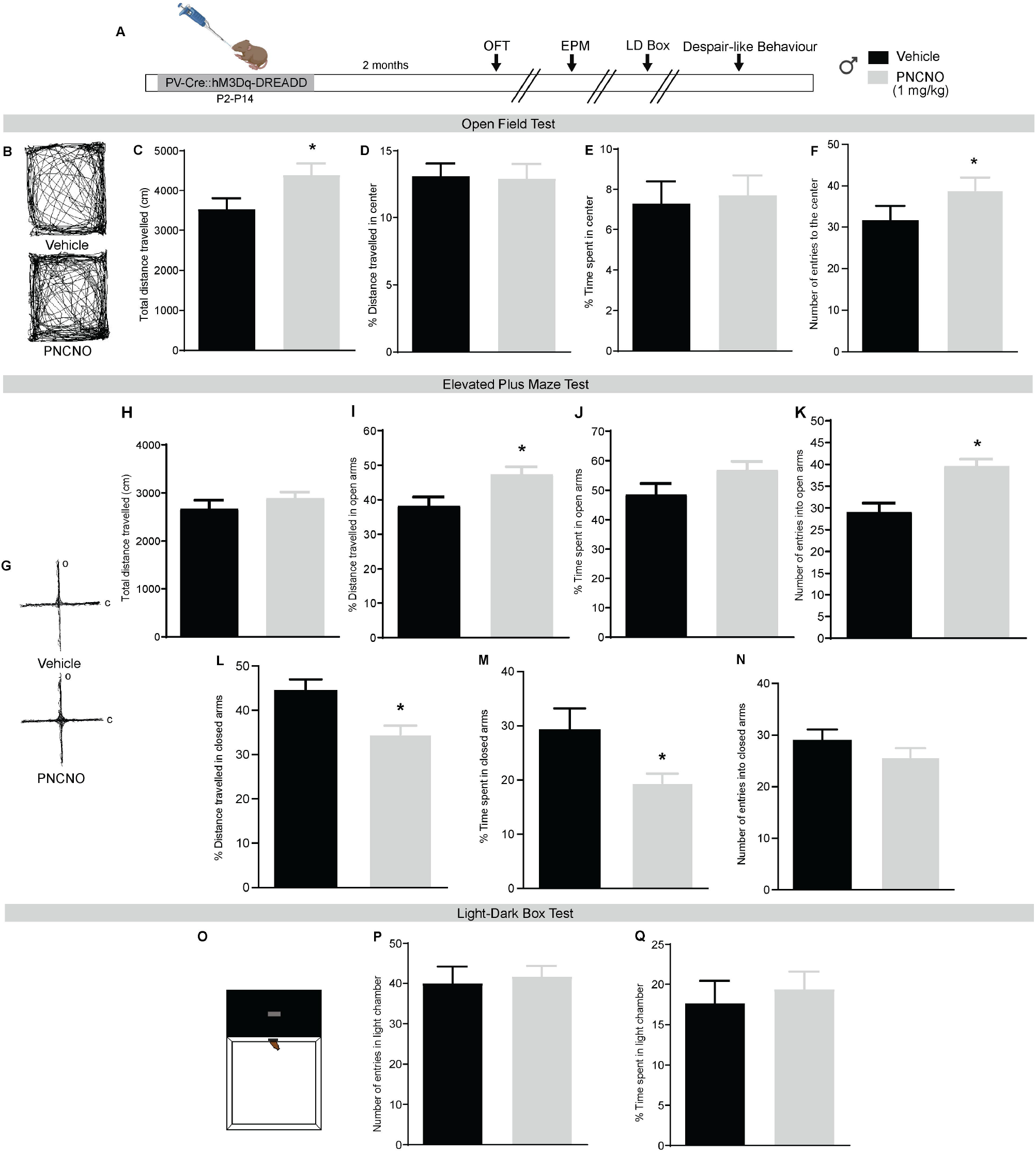
Chronic chemogenetic activation of PV-positive inhibitory interneurons during postnatal life evokes persistent changes in anxiety-like behavior in adult male mice in a task-specific manner (A) Shown is a schematic representation of the treatment paradigm of bigenic PV Cre::hM3Dq-DREADD male mouse pups with vehicle or CNO (1 mg/kg) from P2 - P14 to evoke hM3Dq-DREADD-mediated activation of PV-positive inhibitory interneurons, followed by profiling for anxiety-like behavior on the open field test (OFT), elevated plus maze (EPM), light-dark box test (LD box), followed by behavioral tests for despair-like behavior, with a 7-10 day washout period between all behavioral tests. (B) Shown are representative tracks in the OFT from vehicle and PNCNO-treated male mice. Bar graphs represent the total distance traveled (C), percent distance traveled in the center (D), percent time in the center (E), and number of entries to the center (F) of the OFT arena. (G) Shown are representative tracks in the EPM from vehicle and PNCNO-treated male mice. Bar graphs represent the total distance traveled (H), percent distance traveled in the open arms (I), percent time spent in the open arms (J), number of entries to the open arms (K), percent distance traveled in closed arms (L), percent time spent in the closed arms (M), and number of entries to the closed arms (N) of the EPM. (O) Shown is a representative image of a LD box. Bar graphs represent the number of entries into the light chamber (P), and percent time spent in the light chamber (Q). Results are expressed as the mean + S.E.M. (\**p* < 0.05 as compared to their vehicle treated controls, n = 17 - 23 per group, Student’s *t-*test).

We then performed behavioral analysis for vehicle and PNCNO-treated bigenic PV-Cre::hM3Dq-DREADD females in adulthood on the OFT, EPM and LD box (Figure 4A). In the OFT, we noted no significant differences between the treatment groups on any of the measures analyzed, namely the total distance travelled in the arena (Figure 4C), the percent distance and percent time in the center of the arena (Figure 4D, E) or in the number of entries to the center of the arena (Figure 4F). In the EPM test, PNCNO-treated female mice exhibited a significant increase in total distance travelled in the maze (Figure 4H, *p* = 0.01) as compared to the vehicle-treated cohort, but did not differ on other measures namely the percent distance or time in the open arms (Figure 4I, J), percent distance or time in the closed arms (Figure 4L, M), or the number of entries to the open or closed arms (Figure 4K, N). In the LD Box, PNCNO-treated female mice exhibited a significant increase in the percent time spent in the light chamber (Figure 4Q, *p* = 0.02), but did not show any difference from controls for the number of entries to the light chamber (Figure 4P). Taken together, these results indicate that a history of chemogenetic activation of PV-positive neurons during postnatal life (P2 - 14) is associated with a decline in anxiety-like behavioral responses in adulthood in males and females, which are noted in a task-specific manner.

**Figure 4.**
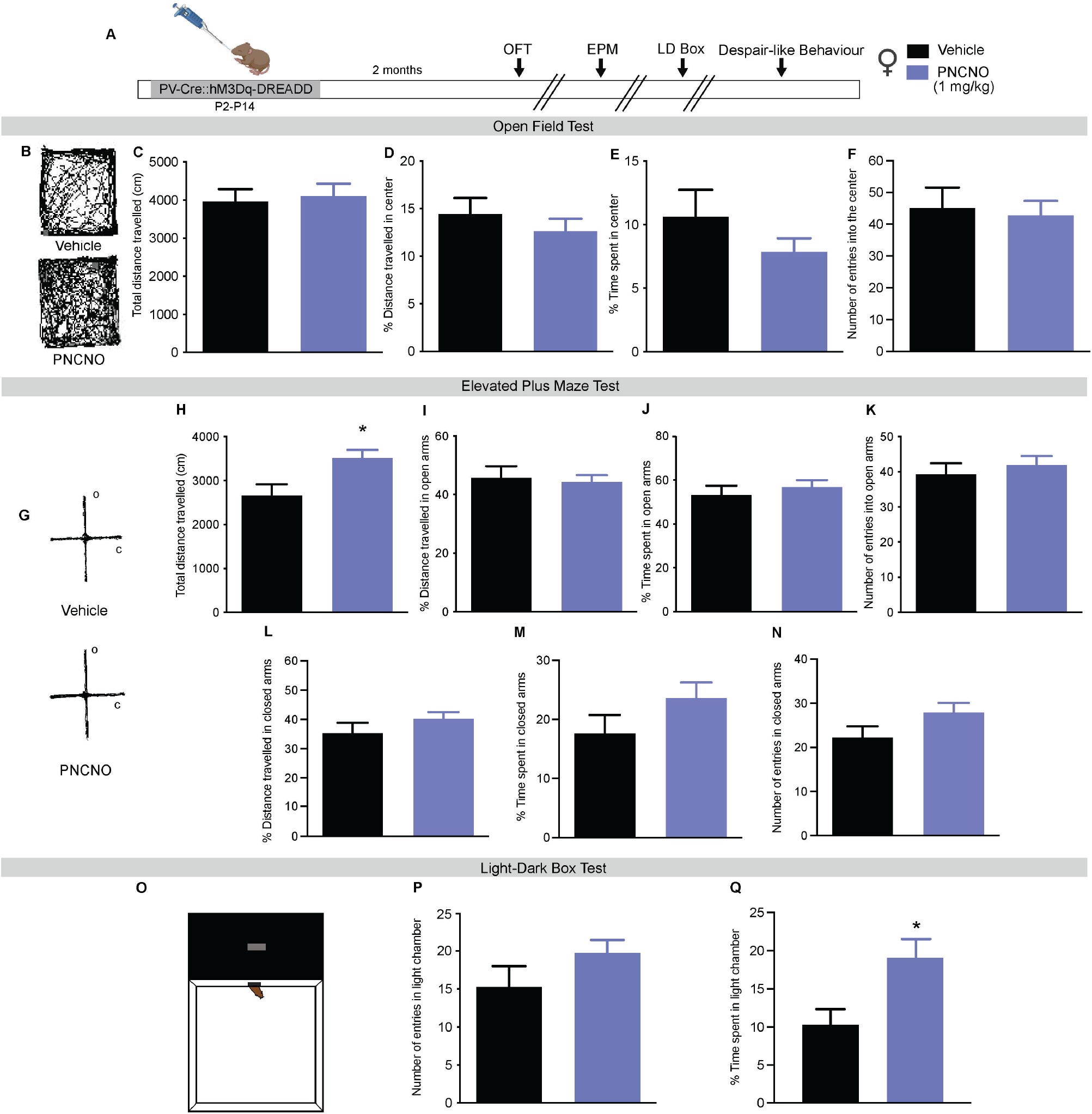
Chronic chemogenetic activation of PV-positive inhibitory interneurons during postnatal life evokes persistent changes in anxiety-like behavior in adult female mice in a task-specific manner (A) Shown is a schematic representation of the treatment paradigm of bigenic PV Cre::hM3Dq-DREADD female mouse pups with vehicle or CNO (1 mg/kg) from P2 - P14 to evoke hM3Dq-DREADD-mediated activation of PV-positive inhibitory interneurons, followed by profiling for anxiety-like behavior on the open field test (OFT), elevated plus maze (EPM), light-dark box test (LD box), followed by behavioral tests for despair-like behavior, with a 7-10 day washout period between all behavioral tests. (B) Shown are representative tracks in the OFT from vehicle and PNCNO-treated female mice. Bar graphs represent the total distance traveled (C), percent distance traveled in the center (D), percent time in the center (E), and number of entries to the center (F) of the OFT arena. (G) Shown are representative tracks in the EPM from vehicle and PNCNO-treated female mice. Bar graphs represent the total distance traveled (H), percent distance traveled in the open arms (I), percent time spent in the open arms (J), number of entries to the open arms (K), percent distance traveled in closed arms (L), percent time spent in the closed arms (M), and number of entries to the closed arms (N) of the EPM. (O) Shown is a representative image of a LD box. Bar graphs represent the number of entries into the light chamber (P), and percent time spent in the light chamber (Q). Results are expressed as the mean + S.E.M. (\**p* < 0.05 as compared to their vehicle treated controls, n = 13 - 16 per group, Student’s *t-*test).

### 3.4 Chronic chemogenetic activation of PV-positive inhibitory interneurons during postnatal life alters despair-like behavior in a task-specific manner in male, but not female mice

We next sought to examine whether PNCNO-treated bigenic PV-Cre::hM3Dq-DREADD males exhibit any changes in despair-like behavior on the TST and FST (Figure 5A). In the TST, while we did not observe any change in percent immobility in the PNCNO-treated males as compared to their vehicle-treated controls (Figure 5B), we did note a trend towards a significant decline in the number of immobility events (Figure 5C, *p* = 0.082). On the FST, PNCNO-treated males showed a significant decline in percent immobility (Figure 5D, *p* = 0.018) as compared to their vehicle-treated controls. We did not observe any change in the number of immobility events in the FST (Figure 5E) between the treatment groups. We then addressed whether PNCNO-treated bigenic PV-Cre::hM3Dq-DREADD females show any changes in despair-like behavior on the TST and FST (Figure 6A). We noted no significant difference between the PNCNO-treated bigenic PV-Cre::hM3Dq-DREADD females and their vehicle-treated controls either for the percent immobility noted in the TST (Figure 6B) or FST (Figure 6D), or for the number of immobility events in the TST (Figure 6C) or FST (Figure 6E). Collectively, these results indicate that a history of chemogenetic activation of PV-positive neurons during postnatal life (P2 - 14) does not appear to significantly influence despair-like behavior in female mice in adulthood, whereas the results suggest subtle changes indicative of reduced-despair-like behavior in adult PNCNO-treated males that are noted in a task-specific manner.

**Figure 5.**
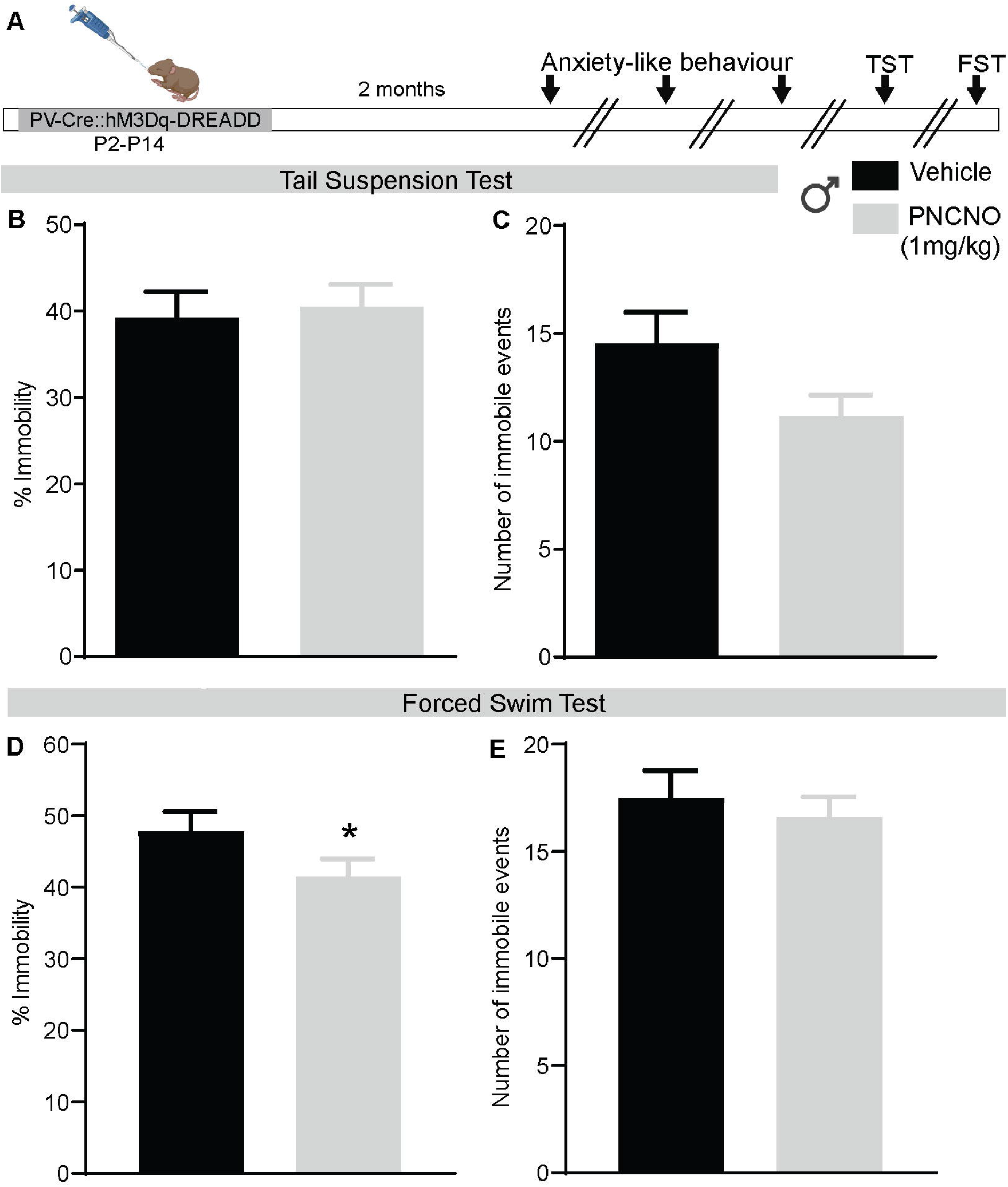
Chronic chemogenetic activation of PV-positive inhibitory interneurons during postnatal life alters despair-like behavior in a task-specific manner in male mice (A) Shown is a schematic representation of the treatment paradigm of bigenic PV Cre::hM3Dq-DREADD male mouse pups with vehicle or CNO (1 mg/kg) from P2 - P14 to evoke hM3Dq-DREADD-mediated activation of PV-positive inhibitory interneurons. Vehicle and PNCNO-treated adult male mice were subjected to a battery of behavioral tasks to profile anxiety-like behavior, followed by profiling for despair-like behavior on the tail suspension test (TST) and the forced swim test (FST), with a 7 - 10 day washout period between behavioral tests. Bar graphs represent the percent immobility (B) and the number of immobile events (C) in the TST. Bar graphs represent the percent immobility (D) and the number of immobile events (E) in the FST. Results are expressed as the mean ± S.E.M. (\**p* < 0.05 as compared to vehicle treated controls, n = 17 - 23 per group, Student’s *t-*test).

**Figure 6.**
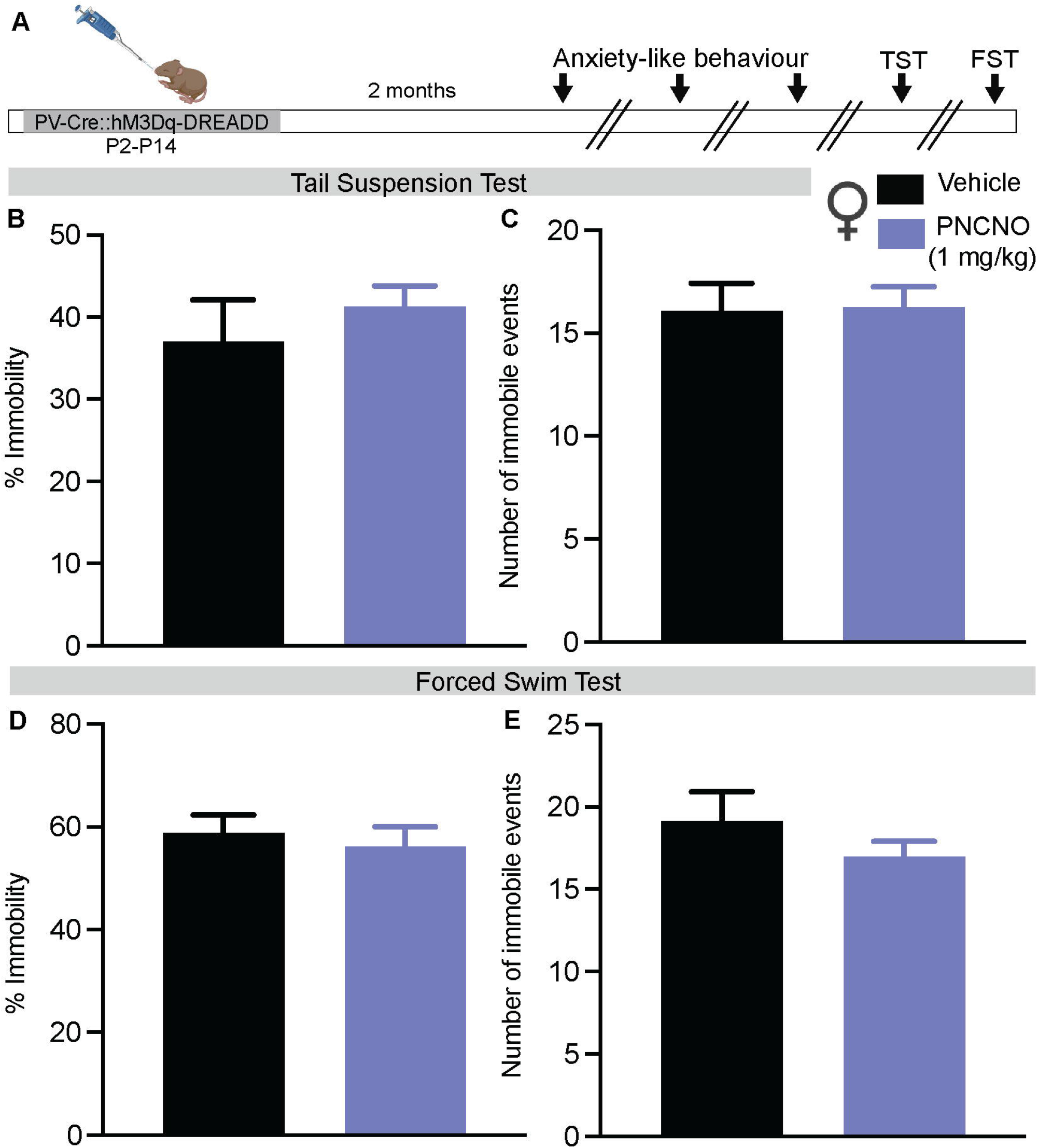
Chronic chemogenetic activation of PV-positive inhibitory interneurons during postnatal life does not alter despair-like behavior in female mice (A) Shown is a schematic representation of the treatment paradigm of bigenic PV Cre::hM3Dq-DREADD female mouse pups with vehicle or CNO (1 mg/kg) from P2 - P14 to evoke hM3Dq-DREADD-mediated activation of PV-positive inhibitory interneurons. Vehicle and PNCNO-treated adult female mice were subjected to a battery of behavioral tasks to profile anxiety-like behavior, followed by profiling for despair-like behavior on the tail suspension test (TST) and the forced swim test (FST), with a 7 - 10 day washout period between behavioral tests. Bar graphs represent the percent immobility (B) and the number of immobile events (C) in the TST. Bar graphs represent the percent immobility (D) and the number of immobile events (E) in the FST. Results are expressed as the mean ± S.E.M. (n = 13 - 16 per group).

## 4 Discussion

Our findings reveal that while chronic chemogenetic activation of PV-positive inhibitory interneurons during the postnatal developmental window does not disrupt sensory milestones or growth, it results in anxiolytic and antidepressant-like behavioral changes in adulthood, which are dependent on biological sex and are observed in a behavioral task-specific fashion.

Prior studies have primarily focused on addressing the role of chemogenetically perturbations of PV-positive interneurons in either juvenile life or adulthood (Zou *et al*., 2016; Chen *et al*., 2018; Page *et al*., 2019; Nawreen *et al*., 2020; Xiao *et al*., 2021). Chronic chemogenetic activation of PFC PV-positive interneurons in adulthood enhances anxiety-like behavioral responses in female mice (Page *et al*., 2019), which is speculated to contribute to the ‘hypofrontality’ noted following chronic stress (Perova *et al*., 2015; Park *et al*., 2016; Page and Coutellier, 2019). Chemogenetic inhibition of PFC PV-positive interneurons enhances active coping behaviors in male mice subjected to chronic stress, offsets stress-evoked physiological consequences, enhances synaptic plasticity and evokes antidepressant-like behavioral effects (Nawreen *et al*., 2020; Fogaça *et al*., 2021). Chemogenetic activation or inhibition of PV-positive interneurons within the hippocampal DG subfield evokes anxiolysis on the EPM in male mice and decreases social interaction respectively (Zou *et al*., 2016; Medrihan *et al*., 2020). Chemogenetic manipulations of PV-positive interneurons in the juvenile window, particularly in the context of animal models of schizophrenia-like behavior indicate an attenuation of the synaptic and behavioral deficits noted in genetic and pharmacological mouse models of schizophrenia (Mukherjee *et al*., 2019; Nieves *et al*., 2020; Huang *et al*., 2021). Selective optogenetic silencing of PV-positive interneurons in the basolateral amygdala in the juvenile window normalizes aberrant fear conditioning responses that arise in an animal models of early adversity (Nieves *et al*., 2020). These chemogenetic and optogenetic studies targeting PV-positive inhibitory interneurons in specific circuits in juvenile and adult animals, highlight an important role in modulation of anxio-depressive behaviors. It also underscores the importance of addressing the role of this subclass of inhibitory interneurons in critical early postnatal temporal windows (Medendorp *et al*., 2021), which are particularly important given the large body of evidence indicating that PV-interneuron number and function is perturbed in both models of early adversity and neurodevelopmental disorders (Rodriguez *et al*., 2018; Ellis and Honeycutt, 2021).

Our results reveal that PNCNO-administered PV-Cre::hM3Dq-DREADD mice appear to exhibit anxiolytic behavioral changes, that are noted in a task-specific and sex-dependent manner. Whilst, PNCNO-treated male mice showed reduced anxiety-like behavior in the EPM and OFT, but not on the LD-box, PNCNO-treated female mice exhibited anxiolytic behavioral changes on the EPM and LD-box, but not the OFT. It is interesting in this regard that on the EPM, which is the most extensively used behavioral test for examining anxiety-like behavior and testing anxiolytic compounds (Carobrez and Bertoglio, 2005; Crawley and Bailey, 2009; Bourin, 2015; Kirlic *et al*., 2017), we note robust anxiolytic responses in adult male, but not female, PNCNO-treated mice. This was also apparent with regards to despair-like behavior on the TST and FST, with PNCNO-treated male, but not female, mice exhibiting a decline in despair-like behavior. This raises the intriguing possibility that postnatal chemogenetic activation of PV-positive interneurons could differentially impact neurocircuits that regulate approach-avoidance behavior and active/passive coping strategies which may underlie the heterogeneity noted in behavioral responses, with a strong influence of biological sex as a key variable (Goodwill *et al*., 2018; Gildawie *et al*., 2020; Ellis and Honeycutt, 2021). Given the differences in the age of onset of treatment, timing, duration and mode of chemogenetic activation, neurocircuit in which PV-positive inhibitory interneurons were targeted, as well as specific behavioral tests performed, it is difficult to directly draw comparisons across our findings and prior studies (Zou *et al*., 2016; Mukherjee *et al*., 2019; Nawreen *et al*., 2020; Huang *et al*., 2021). Nevertheless, collectively these observations suggest that chemogenetically targeting PV-positive interneurons in either postnatal, juvenile or adult windows can influence anxio-depressive behaviors, in keeping with the substantial preclinical and clinical data that implicate this class of neurons in the pathophysiology of neuropsychiatric and neurodevelopmental disorders (Marín, 2012; Dehorter *et al*., 2017).

Prior evidence suggests an interplay between a distributed network of neural circuits in the regulation of anxiety-like behavior, including but not restricted to the PFC, ventral hippocampus, amygdala, nucleus accumbens, septum, hypothalamus and the bed nucleus of the stria terminalis (BNST) (Adhikari, 2014; Duval *et al*., 2015). For example, in the ventral hippocampus, PV-positive interneurons drive theta oscillations, and synchrony of theta oscillations between the ventral hippocampus and mPFC is associated with enhanced exploration in anxiety-like behavioral tests (Adhikari *et al*., 2010, 2011; Amilhon *et al*., 2015). In addition, PV-positive interneurons in the nucleus accumbens and BNST are known to modulate avoidance behavior (Xiao *et al*., 2021). In our experiments, we used a broad PV-Cre driver (Hippenmeyer *et al*., 2005) that would result in the expression of the hM3Dq-DREADD in multiple brain regions, and chemogenetic modulation of PV-positive interneurons during postnatal temporal windows could have implications for the fine-tuning of E/I balance in limbic neurocircuits that influence anxio-depressive behaviors (Ferguson and Gao, 2018; Reh *et al*., 2020). One of the caveats of our study is that rather than a targeted strategy impacting PV-positive interneurons only within a specific microcircuit, we have used a broad PV-Cre driver which would target PV-positive interneurons across diverse neurocircuits (L. Luo *et al*., 2020). However, it is noteworthy that several environmental, pharmacological and genetic perturbation models of neurodevelopmental disorders are reported to impact PV-positive interneuron structure and function across multiple networks (Smith-Hicks, 2013; Goodwill *et al*., 2018; Todorović *et al*., 2019; Soares *et al*., 2020; Aksic *et al*., 2021; Bueno-Fernandez *et al*., 2021; Klimczak *et al*., 2021; Mukhopadhyay *et al*., 2021; Woodward and Coutellier, 2021). In this regard, our approach to chemogenetically manipulate PV-positive interneurons during postnatal development across diverse circuits bears relevance to both models of early adversity and neurodevelopmental disorders, which impact PV-positive interneurons across several limbic brain regions (Ruden *et al*., 2021). Across many of these models a common overarching theme is the emergence of a dysregulated E/I balance (Ferguson and Gao, 2018; Sohal and Rubenstein, 2019). This suggests that chemogenetically perturbing select neuronal populations during these key temporal windows could also impact the establishment of an E/I balance, associated with a programming of aberrant emotionality (Page and Coutellier, 2019; Pati *et al*., 2020; Z. Y. Luo *et al*., 2020).

Previous work from our group has shown that postnatal chronic chemogenetic activation of forebrain excitatory neurons disrupts glutamate/GABA metabolism and programs long-lasting decreases in hippocampal neuronal activity, along with enhancing anxiety and despair-like behavior and perturbing sensorimotor gating (Pati *et al*., 2020). Further, a recent report indicates that postnatal chemogenetic activation of Emx1-positive forebrain neurons led to a reduction of social behavior and enhanced grooming behavior, associated with a decreased cortical neuronal excitability and enhanced E/I ratio (Medendorp *et al*., 2021). In contrast, a chemogenetic inhibition of forebrain excitatory neurons during the postnatal window did not influence adult anxiety and despair-like behavior (Tiwari *et al*., 2022). In this regard, our findings that postnatal chemogenetic activation of PV-positive inhibitory interneurons can program a reduction in anxiety and despair-like behavior, suggests that targeting specific neuronal subpopulations in key limbic microcircuits during this window can program differing behavioral outcomes for mood-related behaviors. To the best of our knowledge few studies have addressed whether manipulation of neuronal activity within PV-positive interneurons during the early postnatal temporal window impacts the shaping of cortical microcircuits and emergence of mood-related behavior (Wong *et al*., 2018; Huang *et al*., 2021; Medendorp *et al*., 2021). In a prior study, bioluminescent optogenetic approaches to enhance activity in cortical PV-positive interneurons from P4 – P14, did not appear to change social approach behavior (Medendorp *et al*., 2021). Previous work suggests that chemogenetically enhancing pyramidal neuron activity during postnatal windows of PV-positive interneuron cell death can dampen interneuron cell loss and increase PV-positive interneuron density, with key controls indicating that this is not an off-target effect of the DREADD agonist, CNO (Wong *et al*., 2018). This raises the speculative possibility that chemogenetic strategies that impinge on altering activity of specific neuronal populations during critical developmental windows could via a disruption of the normal developmental trajectory of E/I balance and the sculpting of microcircuits, influence the programming of adult trait anxiety states (Soiza-Reilly *et al*., 2019; Page and Coutellier, 2019; Sohal and Rubenstein, 2019; Pati *et al*., 2020; Teissier *et al*., 2020; Adjimann *et al*., 2021; Medendorp *et al*., 2021; Tiwari *et al*., 2021). The short and long term consequences of chemogenetic manipulation of PV-positive interneurons on microcircuit reorganization and eventual network rhythmogenesis has been relatively less well studied (Wong *et al*., 2018; Huang *et al*., 2021; Rogers *et al*., 2021). A recent study highlights paradoxical effects of CNO-mediated DREADD activation on population firing rates of inhibitory and excitatory neurons, as well as changes in long term plasticity, providing ample caution whilst considering the mechanisms targeted by DREADD manipulations (Rogers *et al*., 2021). The temporal windows used in multiple early adversity models overlaps with the time-window used for chemogenetic manipulations of PV-positive interneurons (Katahira *et al*., 2018; Gomes *et al*., 2019; Reh *et al*., 2020; Tiwari *et al*., 2021), and this opens up the possibility that such postnatal modulation of PV-positive interneuron activity during critical periods could directly impinge on the manner in which local microcircuit rhythmogenesis is shaped (Sohal *et al*., 2009; Moran *et al*., 2017; Murthy *et al*., 2019), with biological sex playing a key role given that PV interneurons also express sex hormone receptors (Wu *et al*., 2014). A recent review raises the intriguing possibility that “estrogens may serve a protective role” resulting in the “blunting” of some of the impact of early adversity in females (Ellis and Honeycutt, 2021; Woodward and Coutellier, 2021). Our results further highlight that biological sex is a key variable to keep in mind when assessing the impact of PV-positive interneuron manipulations in the shaping of adult emotionality.

Collectively, our findings indicate that enhanced activation of PV-positive inhibitory interneurons during early postnatal developmental windows can shape the emergence of anxiety and despair-like behavioral traits in adulthood, with a reduction noted in anxio-depressive behaviors in a sex and behavioral task-dependent fashion. These observations motivate future investigation to directly address how the modulation of PV-positive interneuron activity in this critical temporal window may influence microcircuit development and rhythmogenesis, the formation and maintenance of PNNs, and the sex-dependent and circuit-specific contribution of specific PV interneuron populations in regulation of trait anxiety and despair-like behavioral states.

## 6 Author Contributions

Author contributions: T.B. and V.A.V. designed research; T.B., S.P., and P.T. performed research; T.B., and P.T. analyzed data; T.B. and V.A.V. wrote the paper.

## 8 Acknowledgments

We thank Dr. Shital Suryavanshi, and the animal house staff at the Tata Institute of Fundamental Research, Mumbai, for technical assistance. We thank Prof. Neeraj Jain, National Brain Research Centre, Manesar for the sharing of specific reagents.

## 9 Contribution to the field statement

The pathogenesis of neurodevelopmental and neuropsychiatric disorders is thought to involve alterations in the number and the function of parvalbumin-positive inhibitory interneurons that can disrupt the balance of excitation and inhibition in neurocircuits, setting in motion a trajectory for vulnerability to psychopathology. PV-positive interneurons are particularly sensitive to environmental, pharmacological and genetic perturbations during critical early postnatal developmental windows. Here, we sought to address whether chemogenetic activation of PV-positive interneurons during postnatal life shapes the emergence of anxio-depressive behaviors in adulthood. We find that chronic chemogenetic activation of PV-positive inhibitory interneurons during postnatal life leads to a decline in anxiety and despair-like behavior in adulthood, in a behavioral task-dependent and sex-specific manner. These results, along with studies examining the influence of chemogenetic perturbations of PV-positive interneurons in the juvenile and adult window, indicate that modulative neuronal activity in this subclass of inhibitory interneurons can influence anxio-depressive behavioral states.

